# Complement modulation synergizes with therapeutic hypothermia in a rat model of neonatal HIE

**DOI:** 10.64898/2026.04.07.717097

**Authors:** Angela Saadat, Haree Pallera, Frank Lattanzio, Dania Jacubovich, Stephanie Newman, Meghana Kunam, Andreea Necula, Aliyah Mohammed, Tushar Shah

## Abstract

**Background:** Neurodevelopmental impairment remains common in neonatal hypoxic–ischemic encephalopathy (HIE) despite treatment with the standard of care, therapeutic hypothermia (TH). The complement response activates at reperfusion and is known to exacerbate neuroinflammation and injury, though its full role and interaction with hypothermia are incompletely defined. We hypothesized that modulating the complement response could improve structural and functional outcomes in HIE, and tested a novel complement therapy (CT), consisting of C3a peptides and the C5a-receptor antagonist PMX205, as both a stand-alone treatment and as an adjuvant to TH.

**Methods:** Wistar rat pups were randomized to the following treatment groups: Sham (uninjured control), NT (uninjured, normothermia/not treated control), or injured and treated with either TH, CT, or CT+TH. At term-equivalence, mild-moderate hypoxic-ischemic injury was induced by Vannucci’s method. To capture the short and long-term effects of the treatments, cohorts were harvested 3 or 66-72 days post-injury, respectively. Cerebral injury was measured by quantifying levels of inflammatory markers and cerebral tissue loss, and functional outcomes were assessed in a series of behavioral tests. The data were stratified to detect sexual dimorphisms.

**Results:** CT and TH treatments demonstrated test and sex-dependent differences in improvement compared to untreated, injured rats. In male rats, TH treatment worsened long-term hippocampal and thalamic brain injury and functional measures of ataxia and attention. CT-treatment worsened long-term thalamic loss in females. Combining the two treatments (CT+TH) demonstrated additive improvement in both sexes, including short and long-term cortical loss and ataxia.

**Conclusions:** Complement modulation enhances the neuroprotective effects of TH after neonatal hypoxic–ischemic injury, with sex-specific effects on inflammation and behavior. Combining complement modulation with the standard of care often demonstrated synergistic improvement in both sexes, supporting complement-targeted therapy as a promising adjunct to hypothermia in neonatal HIE.

**Graphical abstract.:** 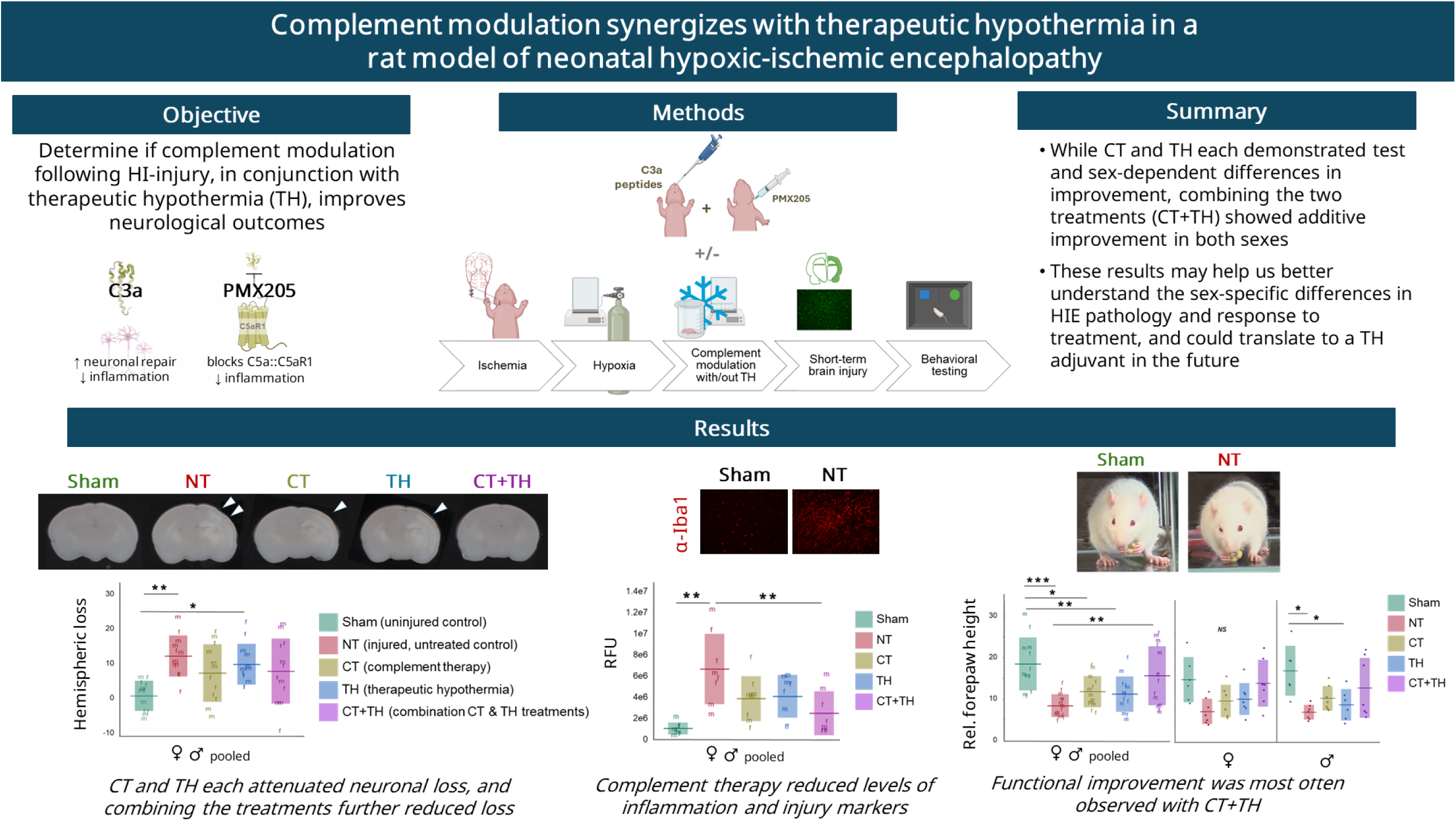

Created with BioRender. Saadat, A. (2026) https://BioRender.com/siwm825.

## Introduction

Neonatal hypoxic-ischemic encephalopathy (HIE) remains a critical global health challenge ^1–4^, with therapeutic hypothermia (TH) serving as the sole evidence-based neuroprotective intervention ^5^. TH reduces mortality and improves early childhood outcomes among survivors, yet significant residual disability persists, and no adjunct treatments to TH have been clinically validated ^6,7^. This therapeutic stagnation underscores the urgent need to unravel HIE’s molecular complexity to identify customized novel, mechanistically driven interventions ^8^.

Recent clinical trials highlight both the limitations of current approaches and the necessity for mechanistic innovation. The HELIX trial ^9^ revealed TH’s inefficacy in low- and middle-income countries (LMICs), where 50% of cooled infants still died or developed disabilities, potentially due to distinct injury patterns from subacute hypoxia in out born populations. Conversely, the HEAL trial ^10^ explored erythropoietin as an adjuvant to TH but faced challenges in demonstrating clear efficacy, reflecting the difficulty of translating preclinical neuroprotection strategies to heterogeneous clinical populations. These trials collectively emphasize that improving outcomes requires moving beyond empiric combination therapies toward interventions grounded in precise pathophysiological insights.

Emerging research on regulated cell death mechanisms—including necroptosis, pyroptosis, and ferroptosis—further illustrates HIE’s biological complexity, suggesting that combinatorial approaches targeting multiple injury phases may be essential ^11,12^. The complement system exemplifies a mechanistically rational target for development of a therapeutic adjunct for TH. While complement activation exacerbates blood-brain barrier disruption and neuroinflammation post-HIE ^13–17^, its dual role in neurodevelopmental processes ^18–22^ necessitates selective modulation. C3a and C5a are central to all complement pathways. C5a is a potent anaphylatoxin and engagement of C5a with its receptor C5aR1 induces microglial activation, neuronal apoptosis and sustained neuroinflammation ^23–25^. HIE increases systemic C5a levels expression of its receptor C5aR1 in the brain ^26^, and inhibiting C5a-C5aR1 engagement with a small molecular inhibitor PMX205 ^27^ improves structural and functional outcomes in HIE ^28^. C3a plays a paradoxical role in neuroinflammation and has anti-inflammatory roles in acute phases of injury, directly countering the effects of C5a ^29^. C3a has been shown to stimulate neurogenesis and neuroplasticity following ischemic brain injury, and elevating C3a levels by the intranasal route decreases neurodegeneration and gliosis ^30–33^.

Integrating mechanistic discoveries with clinically adaptable delivery systems could enable therapies that synergize with TH’s metabolic suppression while preserving neurodevelopmental trajectories. We hypothesized that neurological damage can be mitigated in neonates following hypoxic ischemic insult by concomitantly blocking C5a-C5aR1 interaction and increasing cranial levels of C3a, and that this combination therapy will synergize with TH to improve functional and structural outcomes. Utilizing this targeted, nuanced approach in a rigorous preclinical model that replicate human disease heterogeneity may aid in bridging the gap between bench and bedside ^9^.

## Methods

### Animals

Two independent animal experiments were performed. A short-term cohort was harvested 3 days-post-injury (dpi) to compare injury accumulated in the initial phases of HIE (primary into secondary phases) ^34^, and a long-term cohort at 66-72 dpi to assess injury accumulating into the tertiary injury phase ^34^ (Fig. 1). Throughout, timed-pregnant rat dams (*Rattus norvegicus,* Wistar strain) were procured from Hilltop Lab Animals Inc. (Scottsdale, PA) at embryonic day 19, housed individually, and allowed to deliver spontaneously. At post-natal day 1-3 (P1-3) the pups were pooled, culled, and randomly redistributed among the dams to control for litter effects. The rats were next randomized to treatment groups: uninjured control (Sham), injured and untreated/normothermia control (NT), injured and TH-treated (TH), injured and treated with complement therapies (CT), or injured and treated with both complement and TH treatments (CT+TH), such that each dam was assigned pups of each treatment group. At P21, the long-term cohort pups were randomized into same-sex, different-treatment pairs per cage, using the randomization function in MS Excel, and weaned to Teklad rat chow with Nutra-Gel supplementation. Due to the odd number of animals, the minimum number of cages in each cohort had 3 same-sex rats. Fruity Gems were given post-weaning as enrichment.

**Figure 1.**
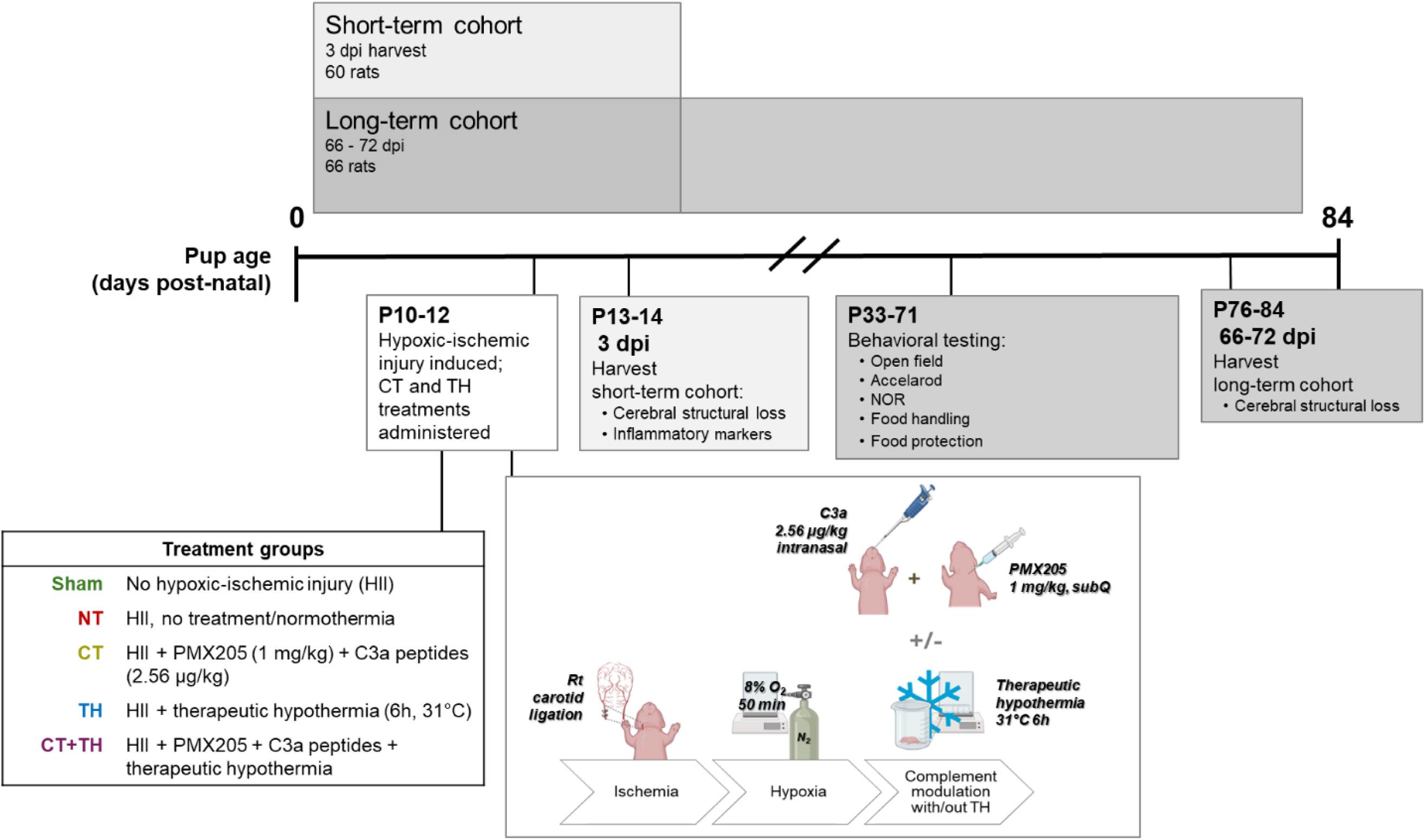
Experimental overview. Two independent animal experiments were performed; one harvested at 3 dpi to compare brain injury accumulated from the initial phases of HIE pathology (light grey box, short-term cohort), the other harvested into adulthood to assess the functional and cumulative effects of primary, secondary, and tertiary phases on brain injury (dark grey box, long-term cohort). In both experiments, rats were randomized to 5 treatment groups: Sham, NT, CT, TH, and CT+TH (defined at lower left). Injury was induced by unilateral carotid ligation and oxygen restriction (Vannucci’s model) and treatments were administered on the day of injury. The long-term cohort animals performed a series of behavioral tests and brains were harvested from both cohorts for histopathological assessments. Created with BioRender. Saadat, A. (2026) https://BioRender.com/siwm825.

The rats were housed in the Virginia Health Sciences at Old Dominion University [VHS at ODU; formerly the Eastern Virginia Medical School (EVMS)] Comparative Medicine facility in a temperature and humidity-controlled room (68-76°F, 30-60%) with a 12-hour diurnal light/dark cycle. All procedures and testing were carried out in this facility or our laboratory in the same building.

126 total rats were used in these experiments. The short-term cohort contained 60 rats total, with 12 rats in each treatment group (Sham, NT, CT, TH and CT+TH) and equal gender distribution. The long-term injury cohort consisted of 66 rats total, with 12 rats in the Sham group (6 per sex), 14 rats in the NT group (7 per sex), 13 rats in the CT group (6 females, 7 males), 13 rats in the TH group (7 females, 6 males), and 14 rats in the CT+TH group (7 per sex). (The 7^th^ CT female and 7^th^ TH male died during injury induction.) The animals were treated in accordance with EVMS IACUC protocol #22-008.

### Hypoxic-ischemic injury

Mild-moderate HIE was induced at P10-12 with Vannucci’s method ^35^ as described previously ^20,36^. Briefly, pups were anesthetized by isoflurane inhalation (3% induction, 1% maintenance) and ischemia was induced by permanent ligation of the right common carotid artery (4-0 silk suture), at two sites 2-5 mm apart. Sham rats received anesthesia, incision, and dissection but no ligation. Incisions were sealed with VetBond and animals were allowed to recover (∼10 min) returning to the dam for 1 hour to rest and feed. For hypoxia, the ligated animals were placed in a custom chamber (BioSpherix) balanced to 8% oxygen with nitrogen gas at 37°C for 60 min (including a 10 min period where the chamber returns to 8% O_2_ following opening to place the animals inside). To control for this time away from the dam, Sham rats were placed in an identical chamber at ambient oxygenic conditions at 37°C.

### Complement treatment

PMX205 trifluoroacetate salt (Cayman Chemical) was reconstituted in 5% ethanol then diluted to 0.25 μg/μL in PBS ^37^. This solution was injected subcutaneously ^37^ into the neck scruff for a total dose of 1 mg/kg. C3a peptides (Complement Technology, Inc.) were serially-diluted to 0.008 μg/μL in PBS and slowly pipetted into each nostril, as by Pekna and colleagues^38^, for a total dose of 2.56 μg/kg. Both drugs were adjusted for variations in pup weight, warmed to ∼37°C, and administered as a one-time dose immediately following recovery from the hypoxia procedure; Sham, NT, and TH animals received vehicle control inoculations.

### Therapeutic hypothermia

After resting with their dam for 1 h post-hypoxia, TH pups were placed in a temperature-controlled chamber (Biospherix) for 6 consecutive hours, inside individual beakers to prevent huddling, where rectal temperatures 31°C +/- 1°C were maintained after the first 15 min. To control for this time away from the dam, rats from the Sham, NT, CT and CT+TH groups were placed in a warmed chamber for a target rectal temperature of 37°C.

### Behavioral tests

Pups in the long-term cohort were evaluated in 4 behavioral tests from ages 3-11 weeks (P25-77). To minimize confounding variables, testing order was mixed, equipment was cleaned between animals with 70% ethanol, and the sexes were tested consecutively each day post-estrous cycle onset.

#### Open field

The rats performed this test at P27-29. Rats were placed alone in the center of a round, plastic table (1.22 m diameter) and allowed to freely explore the field. Tests were filmed with a Canon XA30 camcorder in a room dimmed to encourage exploration ^39^ (∼4 lux). To bracket peak exploration time, the first 1.5 minutes of each video were discarded^40–43^ and the end time trimmed to 8 min. Animal behaviors including the time the rats were mobile were measured with ANY-maze software (Stoelting) in this remaining 6.5 min time window.

#### Food handling

Food handling behaviors spontaneously occur in rodents and are impaired by brain injury ^28,36,44^, and was measured at P35-37 as previously described ^45^. Briefly, rats habituated to the food item, HoneyNut Cheerios, by eating two items a day for the 3 days prior to testing. This item was chosen because all rats in the cohort readily ate them, it was not too heavy for injured rats to manipulate, and it had been used previously ^28,36^. Next the rats were habituated to the apparatus, a 20×20×20 cm glass aquarium then filmed the following day (Canon X16) as they ate at 3 items. Mirrors were placed around the apparatus to increase behavior visibility. 4 rats did not complete their trials in the initial testing day (2 NT, 1 TH, and 1 CT rat).

Images were extracted from the videos based on 1-when the rat’s forelimbs were at their highest position, 2-the food item was >50% of its total size and 3- when at least one profile of the animal was clearly visible. Forelimb height was measured from image stills in ImageJ (Java 1.8.0_172, 64-bit) ^46^ as the length from the 2^nd^- 3rd knuckle of the forepaw to the apparatus floor. To account for variability in their position within the field, these measurements were normalized to body height, which was measured from the highest point of the rat’s midsection to the floor. Relative forelimb height was calculated by dividing forelimb height by body height. Measurements from two images from different food items were averaged for a single, representative mean for each rat.

#### Accelerod

The accelerod test challenges rats to maintain locomotion on a rotating rod that accelerates over time ^47,48^ and training and testing for this was P39-41 and P44-47. The animals were first trained to maintain locomotion on the rotarod (Harvard Apparatus) at a constant speed of 14 rpm for 5 min on days 1 and 2 (non-consecutive), then at 20 rpm for 5 min on day 3. On day 4 the rats were filmed (Canon X16) as they performed 4 trials accelerating from 4-14 rpm over 60 s, with 30 s-1 min to rest between trials. The length of time in each trial that the rats maintained locomotion on the rod was collected from the videos and each rat’s longest time was used for the final analysis. Trials were capped at 280 s to prevent exhaustion and minimize bias, and room lights were dimmed to reduce anxiety and encourage performance.

#### Novel Object Recognition

The novel object recognition (NOR) test was conducted P53-58 over 3 consecutive days. On day 1, the rats were habituated to the apparatus, a 59×48×33 cm black plastic tub, by exploring the empty chamber for 5 min. The following day, rats spent 5 min in the apparatus with two identical objects affixed to opposite corners of the testing tub with double-sided tape. On day 3 rats the apparatus contained one familiar object and one novel object, and time spent engaging with each object were determined offline from videos (Logitech HD and Canon XA30) ANY-maze software (Stoelting), defined as time the animal’s nose was detected within a 2 cm perimeter around each object. The discrimination ratio calculated as: dr = T_novel_ – T_familiar_ / (T_novel_ + T_familiar_)^49^ within the 2 min period (day 3). Room lighting was dimmed to approximately 4 Lux.

#### Food Protection

Food protection ability ^50^ was assessed at P66-71 as we have reported previously.^45^ Briefly, food was mildly restricted to encourage food protection behavior. Restriction began 3 days prior to testing and lasted until the completion of the final test day, for a total of 6 full days. Baseline body mass was established for each rat beforehand, and each day the rats were weighed and given a measured allowance of rat chow. This amount was 6 g for each rat the first day (i.e., 12 g per cage pair), and thereafter was dictated by the percentage of weight lost over each 24-hour period: 6 g was allotted if their weight was equal or less than 10% of their baseline weight, 10 g if between 11-15%, or back to free feeding if their weight was under 15%. *Habituation.* The rats were habituated, in succession, to the food item, accepting the food item from stainless steel forceps, and to the testing apparatus. The food item was given daily for 3 days, in addition to their chow allotment, and was half a Teddy Graham. This item was selected because the rats readily consumed it, its size and weight allowed even the most impaired rats to maneuver with it, and it had been used previously ^45^. Habituation to the apparatus, 58×42×31 cm clear plastic tub, was carried out the day before the first test day. For this, the rats were placed alone in the apparatus and given time to consume their food item for that day, 3-5 min total. *Testing.* Two rats of the same sex, one “robber” and one “victim,” were placed together in the apparatus. The victim rat is given a food item with forceps and tries to simultaneously consume and protecting it from theft. Stolen items were returned to the victim rat to complete the trial if they were 30% or more of the starting size. Each rat completed 3-5 trials each day for 4 total, consecutive days. Only Shams served as robber rats. Day 4 trials were filmed (Canon XA30) aimed at mirrors placed underneath the testing tubs. “Steals” were instances where the robber rat took the food item away from the victim rat and were tallied offline for the final testing day.

### Brain harvest, imaging, and histology

Euthanasia occurred by pentobarbital injection (150 mg/kg, peritoneal, Covetrus) and cardiac puncture and was followed by intracardial perfusion with PBS and 10% formalin. Brains were grossed coronally ∼3 mm from the line of bregma, imaged (Epson Perfection V39 scanner), then paraffin-embedded and sectioned at the VHS Biorepository at ODU (Norfolk VA). Sections were stained with hematoxylin and eosin (H&E) and imaged (Epson Perfection V39 scanner, EVOS M5000 microscope). Individual rat brain images were de-identified and relative structural loss were measured by a blinded researcher in ImageJ (Java 1.8.0_172, 64-bit) ^46^ by tracing the perimeter the entire hemisphere, cortex, hippocampus, and thalamus for each the ischemic (right) and hypoxia-only (left) hemispheres. Relative loss was calculated as 1-(right/left) for each hemisphere and structure.

#### Immunofluorescence and immunohistochemistry

Sections were de-paraffinized (60°C 45 min, xylene washes) and rehydrated with diluted ethanol washes and water. The slides were boiled (10 mM sodium citrate and 0.05% Tween20, pH 6.0) for 20 min. for epitope retrieval, cooled in water, then permeabilized with 0.1% Triton and washed in PBS. TUNEL was probed for on the same brain section as a portion of the anti-GFAP slides; these sections were first blocked and incubated with the TUNEL mixture as instructed by the manufacturer (Roche). All slides were blocked with 5% serum then incubated with primary antibody (Abcam; ab5076, ab4648, ab104224) overnight at 4-8°C then washed. After incubating with secondary antibody (1:500; Invitrogen A11057, A11004), the slides were washed, coverslipped, sealed with nail polish and stored at 4-8°C overnight. Images were captured with a fluorescence microscope (EVOS M5000, Invitrogen; 200x total magnification) where damage was most intense. The images were de-identified and researchers blinded to the treatments measured fluorescence intensity (integrated density) or count (analyze particles) with ImageJ software ^46^.

#### Immunoblot

Protein lysates were prepared from formalin-fixed tissue ^51^. Briefly, 0.05 g of tissue was excised from the ischemic hemisphere, then heated at 100°C for 20 min in extraction buffer (100 mM Tris, 100 mM DTT, and 4% SDS, pH 8.0). Further dissolution of cross-linking followed with 60°C for 2 hours and 40 min of sonication. Lysates were reduced in 20 mM DTT at 37°C for 60 min., then boiled in Laemmli’s buffer (without additional SDS) for 6 min, and stored 2-8°C until use. Samples were re-heated to 37°C for 10 min. in fresh buffer, and 30 µg (1.5 µL) spotted onto PVDF membrane. The blot was blocked with 5% skim milk in TBS, probed with anti-C5aR1 conjugated to Alexa Flour 790 (SC53797) diluted 1:1,000, and washed in TBS with 1% Tween20. Total protein staining was accomplished with Ponceau’s stain (0.5% dye in 0.5% acetic acid), then rinsed with Millipure water. The blots were imaged LI-COR Odyssey M; signal was background-corrected and normalized to total protein values.

### Statistical analyses

Endpoint measurements were performed by researchers blinded to the treatment groups with images deidentified of treatment details; exceptions to this were the NOR, open field, and accelerod endpoints, which were measured by ANY-maze software or the rotarod, respectively. JMP statistical software versions 18-19 was used throughout. Sample sizes were estimated with the sample size explorer tool (power for ANOVA, α<0.05, ß>0.80) using means and within group variances from previous experiments accelerod for the behavioral tests, lesion area for the structural measurements, and anti-Iba1 fluorescence for the inflammatory markers. Datasets were assessed for normality (Shapiro-Wilke’s and Anderson-Darling tests) to determine whether mean or median-based tests were appropriate. In normal datasets, equal variances were determined by p-value<0.05 in 1 or more of O’Brien, Brown-Forsythe, Levene, Bartlett tests. Group differences were determined by Oneway ANOVA if normal, Welch’s ANOVA if normal but variances were unequal variances, or by the Kruskal-Wallis’s test (KW) if the dataset was not normally distributed. Post-hoc Tukey’s tests were used for parametric data sets and Steel-Dwass (SD) for non-parametric datasets. Significance was established as alpha<0.05 throughout. Post-hoc power analyses were performed for all measures, and 0.8 threshold was applied; this and group and individual difference p-values, as well as the specific tests used, are listed in Table 1.

**Table 1.**
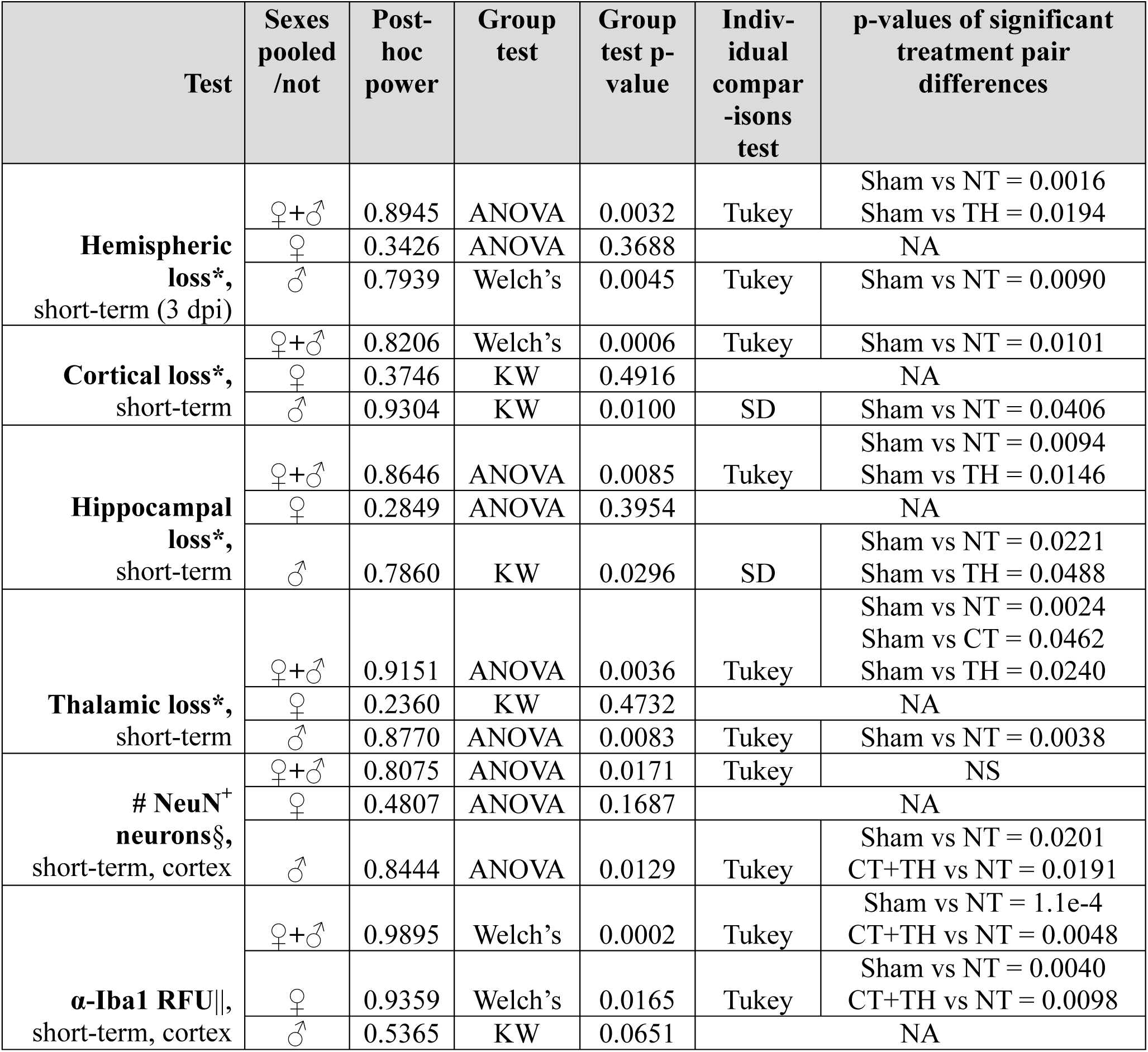
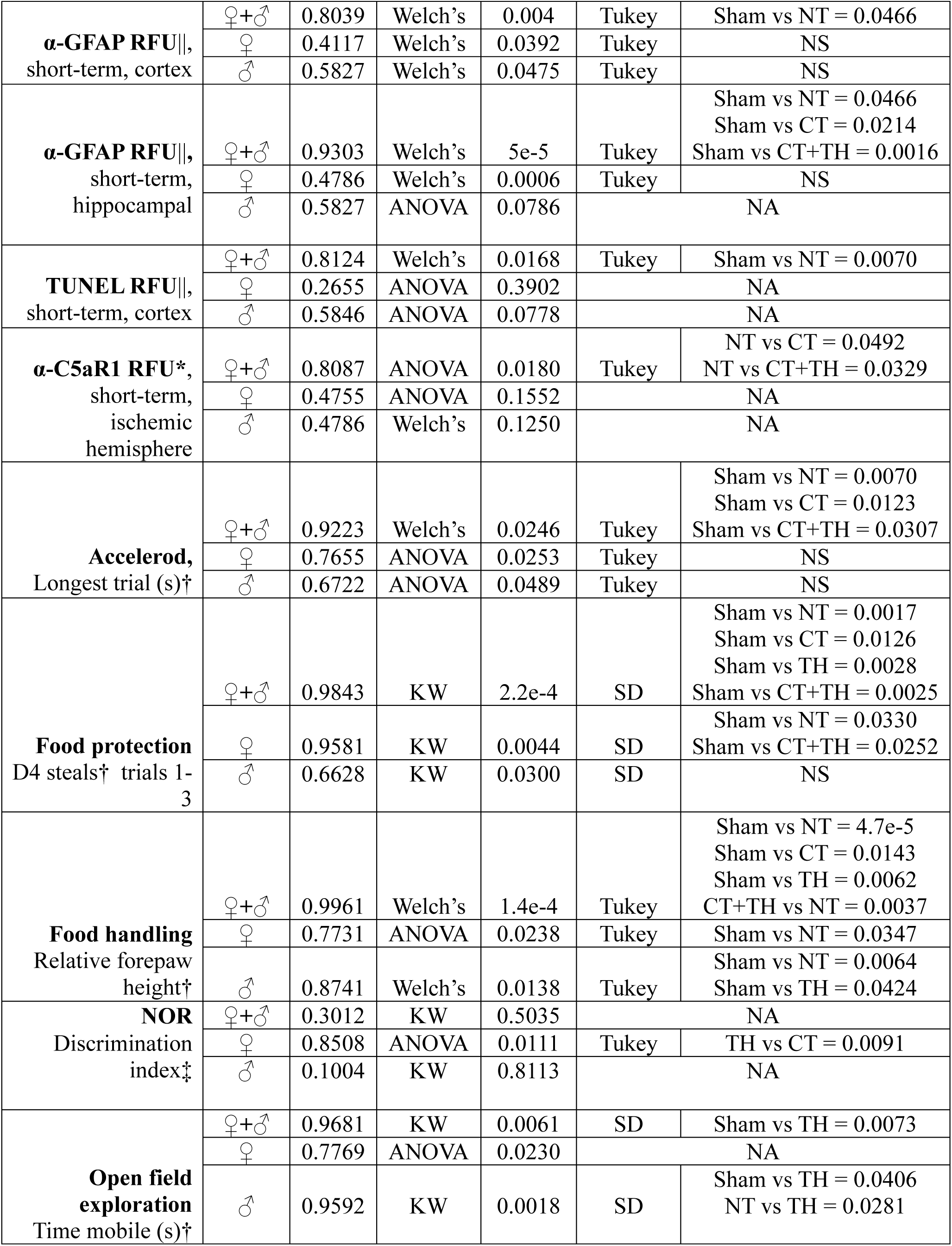
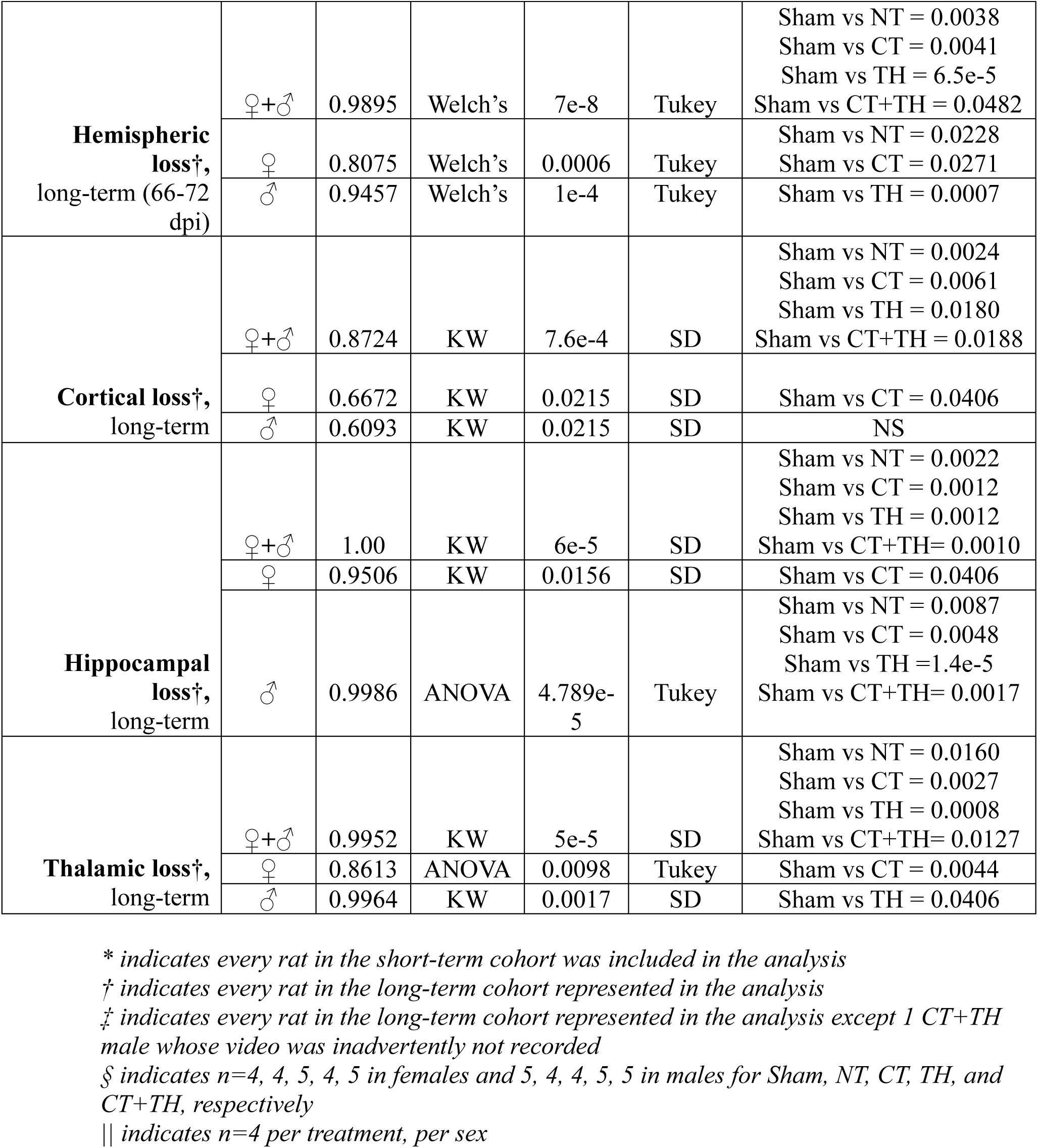
Statistical summary. Listed are each quantitative test, its post-hoc β value, the group test method applied and its p-value, as well as the test used for individual comparisons and the significant p-values. Whether sexes were pooled or stratified (2^nd^ column from left). For the group tests, ANOVA indicates the data were normally distributed, Welch’s indicates the data were normal and variances were equal, and KW indicates one or more treatment groups were not normally distributed. NA indicates a group comparison p-value >0.05, NS indicates ɑ>0.05 by Tukey’s or SD tests.

## 3 Results

### Short-term injury

All injured rats showed increased levels of brain tissue loss compared to Sham (Fig. 2). In the ischemic hemisphere as a whole, NT and TH-treated rats showed the most loss, and CT and CT+TH loss fell between these and Sham, indicating modest improvement (Fig. 2B). In male rats, cortical neuronal density was lowest in NT rats, followed by TH and CT-treated rats, and CT+TH showed the least loss (Fig. 2C). Cortical atrophy was higher in all injured rats, though most extreme in NT rats. In males, NT rats demonstrated the most cortical loss, and only significantly different cortical loss among the injured groups, with CT, TH and CT+TH-treated rat cortical loss falling between NT and Sham measurements (Fig. 2D). NT and TH-treated rats showed the most hippocampal loss, with CT and CT+TH loss being lower but still higher than Sham. In males, thalamic loss was significantly higher in NT rats compared to Sham. TH, CT, and CT+TH-treated thalamic loss were slightly lower than NT and not significantly different from any group. Distinct areas of incomplete liquefaction were also present in 50% of injured rats (Supplementary Fig. 1).

**Figure 2.**
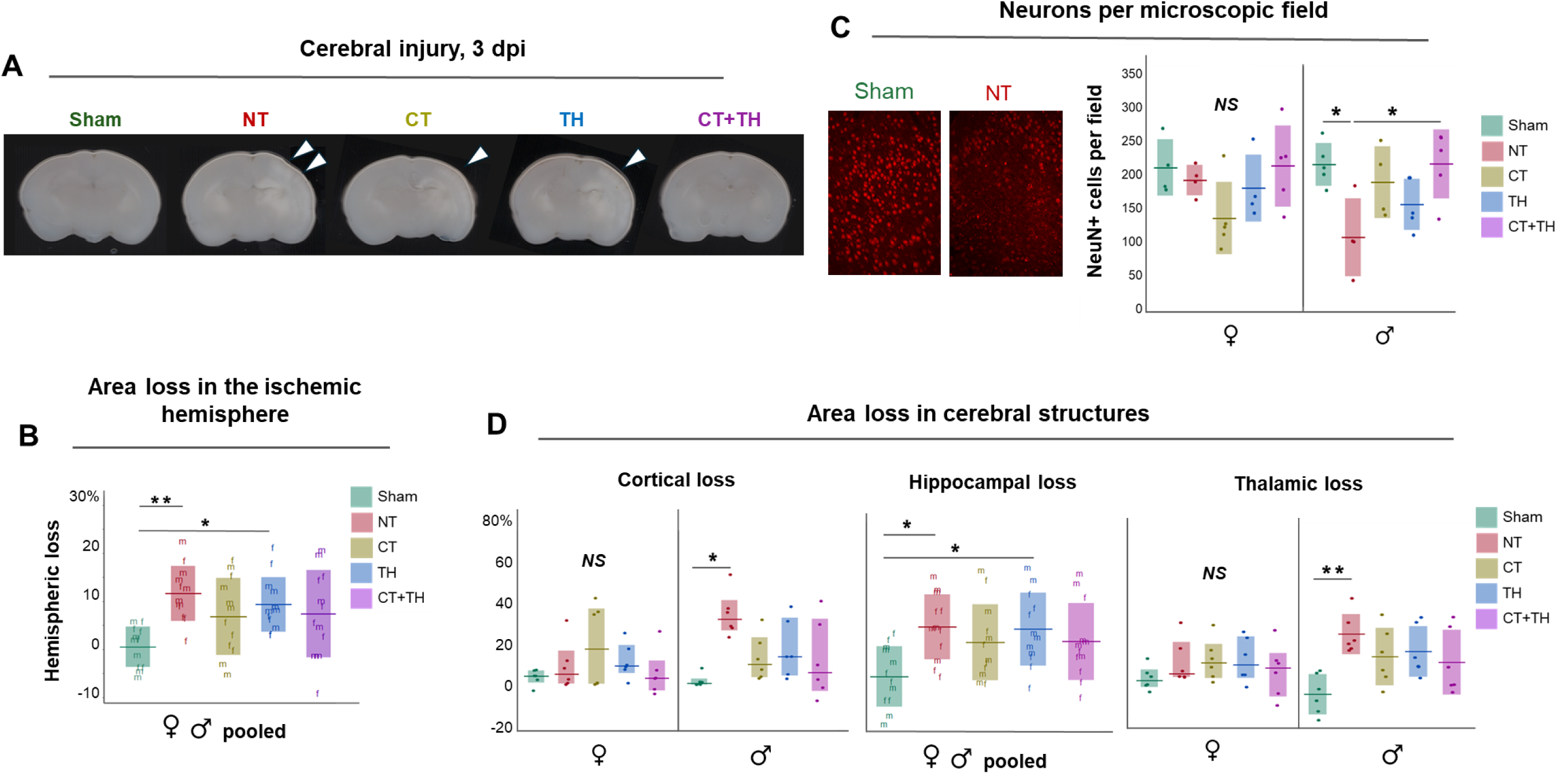
Short-term cerebral injury assessments. **A.** Cerebral lesions were observed as areas of visible tissue reduction in the ischemic hemisphere, varying in severity and common to -2.5 to -5 from the line of bregma. Coronal slices were excised in this region and representative images of each treatment group are shown, positioned so the ischemic hemisphere is on the reader’s right, with white arrows indicating areas of incomplete liquefaction. **B.** Tissue loss in in the ischemic hemisphere is shown as loss in the size (area) of the ischemic hemisphere relative to the left hemisphere, measured from 2D images and calculated as [1- (ischemic hemisphere area/left hemisphere area)]. This was highest in NT rats, followed by TH, CT, CT+TH-treated rats. Sexes were pooled to meet the power threshold. **C.** Neuronal density in the upper cortex was quantified by anti-NeuN immunofluorescence. Representative images (200x total magnification) of the uninjured (Sham) and injured untreated (NT) controls from which the number of NeuN+ neurons within the microscopic field were quantified are shown (at left). Sex-stratification revealed differences were primarily driven by male rats, where density was lowest in NT rats, followed by TH, CT, and significantly improved by CT+TH treatment **D.** Atrophy in the cortex, hippocampus, and thalamus. Cortical and thalamic loss was moderately attenuated in male rats by all treatments compared to NT. Hippocampal injury was highest in NT and TH rats. Horizontal bars represent means (B, C, and hippocampal and male thalamus in D) or medians (both cortical graphs and female thalamic graph in D), and the shaded bars represent standard deviation or interquartile range, respectively. Individual rat values are shown as dots or as “f” and “m” to differentiate female and male values. In C, n=4, 4, 5, 4, 5 in females and 5, 4, 4, 5, 5 in males for Sham, NT, CT, TH, and CT+TH, respectively. In B and D, n=12 per treatment or n=6 per treatment per sex, representing all rats in the short-term cohort. NS indicates significance not reached in post-hoc power and group analyses, *represents p-values from 0.05-0.006, **p-values = 0.005-0.0006, and *** < 0.0005.

### Inflammatory markers

All injured rats showed higher levels of TUNEL signal, but these were highest in NT rats (Fig. 3A), indicating all treatments modestly lowered levels of cortical apoptosis. Iba1 signal differences were primarily by female rats, where NT levels were highest, followed by CT and TH, and levels were significantly improved in CT+TH (Fig. 3B), indicating a synergistic effect of the CT and TH on female microglial activation. GFAP levels in the cortex were higher in all injured rats compared to Sham, though only NT reached significance; in the hippocampus, levels were increased in all injured rats and often higher than NT with treatment, with CT+TH being highest (Fig. 3C). Taken together, all the treatments lowered astrocyte activation in the cortex, but none of the treatments lowered hippocampal levels. Levels of the receptor antagonized by PMX205, C5aR1, were highest in NT rats and modestly lowered by treatments (Fig. 3D).

**Figure 3.**
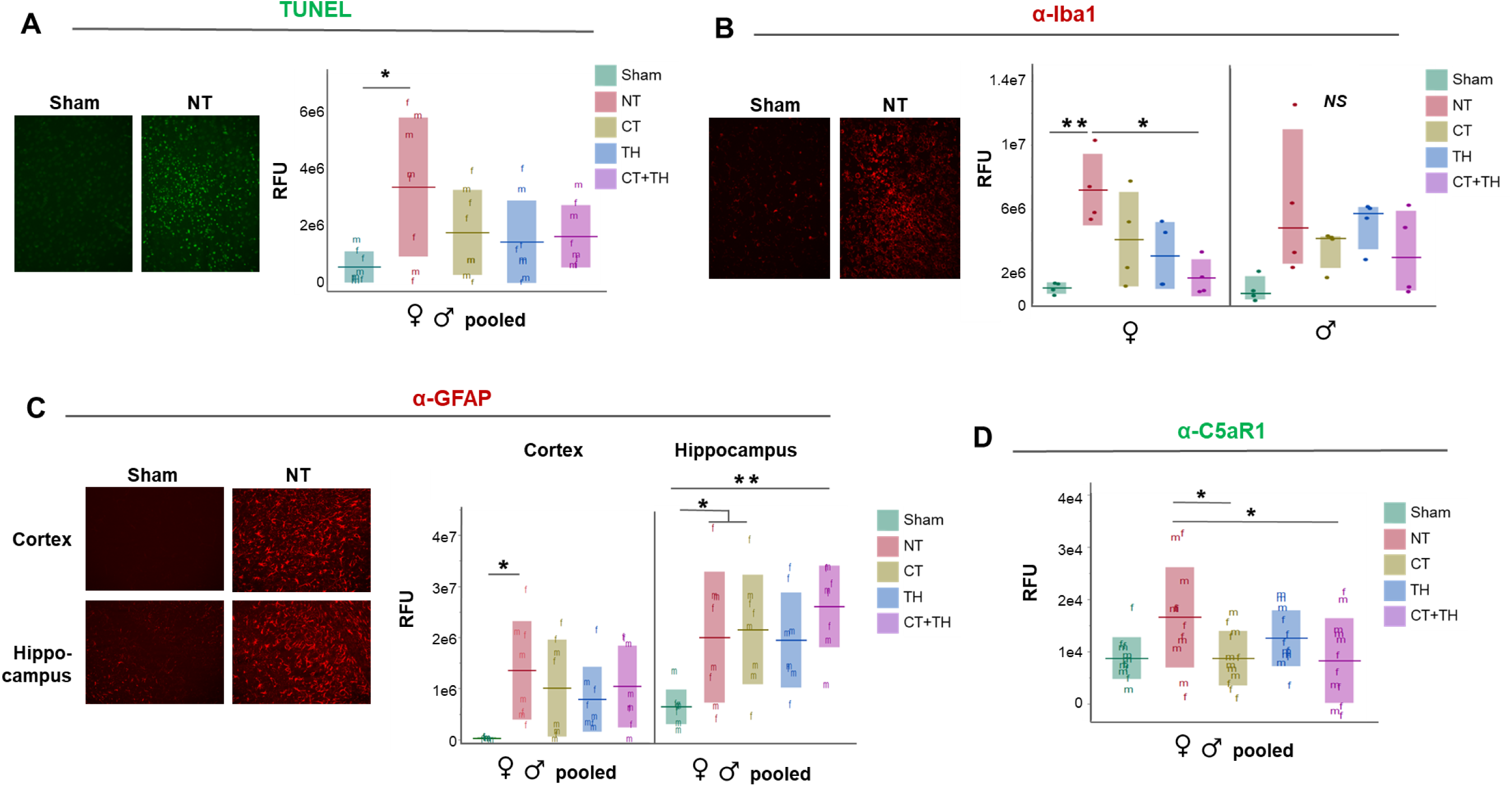
Markers of inflammation in cerebral samples at 3 dpi. Neuroinflammatory markers were measured by densitometry with immunofluorescence (A-C) or immunoblotting (D). **A.** Levels of apoptosis, measured as TUNEL labelling of DNA fragments (green fluorescence), were highest in NT rats and lowered by all treatments. **B.** In female rats, levels of anti-Ibal1 were highest in NT rats, lowered slightly by CT and TH treatments, and further lowered by CT+TH. Female Sham and CT+TH means were different from; male rat data did not reach the power threshold. **C.** Cortical GFAP levels were highest in NT rats, but hippocampal levels were elevated in all also means and standard deviation. **D.** Complement receptor C5aR1 levels were elevated by injury and significantly lowered by CT and CT+TH treatment. Sexes were pooled to meet the power threshold. Shown at left in A-C are representative fluorescence microscopy images (200x) for each uninjured and injured, untreated controls are. In graphs A, C, D, and the female portion of B, the horizontal bars represent means and the shaded bars represent standard deviation; in the male graph of B, these represent median and interquartile range. Sexes were pooled in A, C and D to meet the power threshold. Individual rat values are shown as dots or “f” and “m” to differentiate female and male values. Sample sizes in A-C are n=4 per treatment, per sex; in D in n=12 per treatment group. * represents p-values from 0.05-0.006, ** p-values = 0.005-0.0006, and *** < 0.0005.

Elevated expression of C5aR1 is associated with neuroinflammation and predominantly driven by microglial activation ^52^, and to a lesser extent by neurons ^23,53^, astrocytes, endothelial cells ^54,55^, and infiltrating granulocytes and monocytes ^56,57^. PMX205 and its sister molecule PMX53 ^58^ have been shown to reduce C5aR1 transcripts and expression following ischemic brain injury ^52,53^. Taken together, the increased levels of C5aR1 levels were likely largely due to elevated microglial levels, which were variably attenuated by all treatments (Fig. 3B).

### Long-term functional & injury comparisons

#### Open field

Time mobile during exploration was measured as an indicator of disordered attention and hyperactivity ^59^. Collectively, TH-treated rats spent more time mobile than the rats in all other treatment groups. In males, TH-treated rats spent significantly more time mobile than both Sham and NT rats, 30.6% more time than Sham and 26.1% more time mobile than NT (Fig. 3A).

#### Accelerod

The accelerod test was used to measure balance, coordination, and locomotion deficits from HI-injury. (Fig. 4B). All injured groups showed deficit, though TH-treated rats demonstrated the least impairment, largely driven by three high female values (visible above the standard deviation bar in Fig. 4B), though when stratified by sex, the data not meet power threshold.

**Figure 4.**
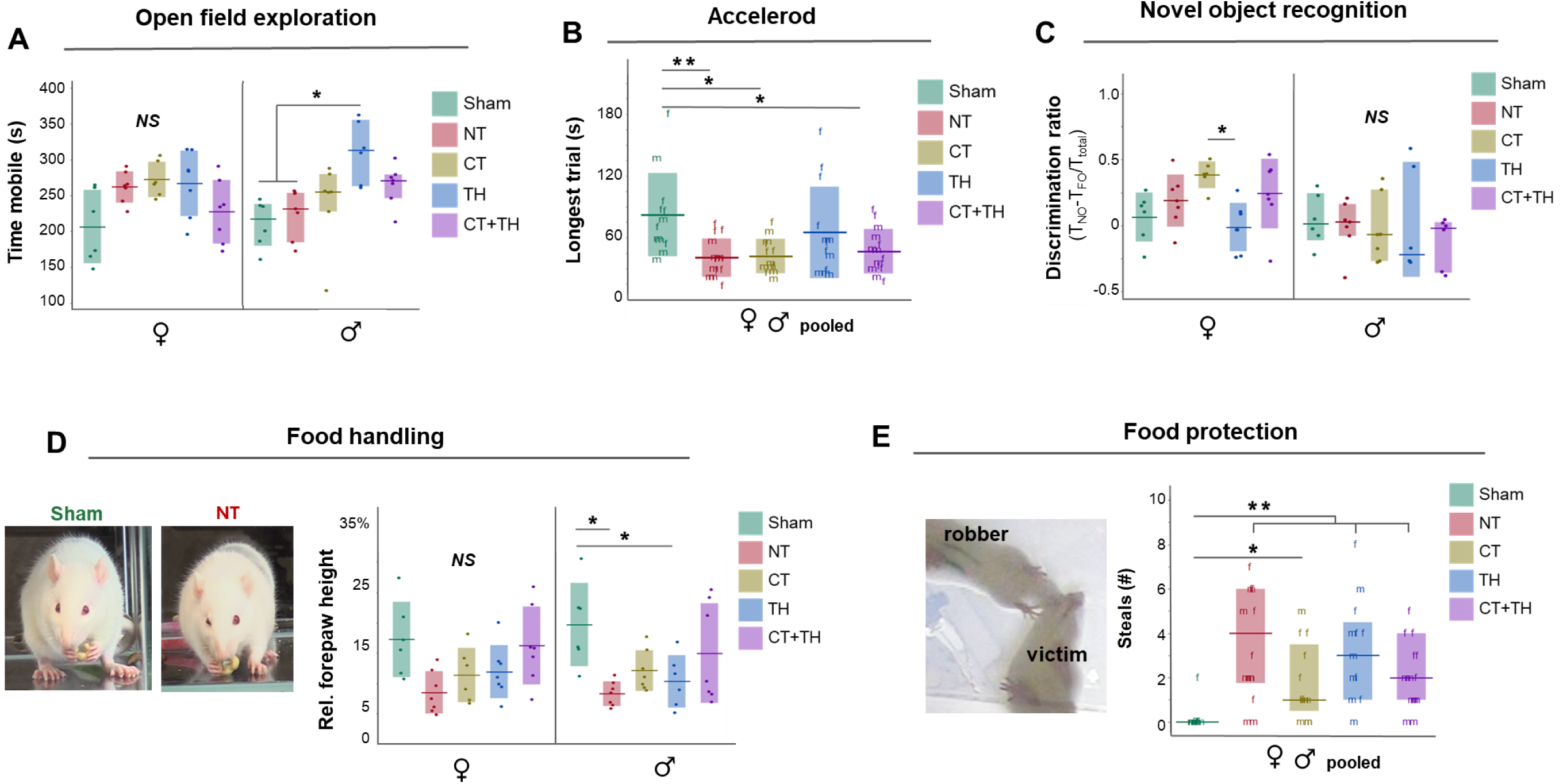
Functional outcomes. **A**. In the Open field test, TH-treated male rats spent more time mobile (in seconds) than Sham and NT rats, while female rats did not demonstrate statistical differences. **B.** In the accelerod test, all injured rats were less able to maintain locomotion on the rod than Sham. When ranked by p-value, NT rats demonstrated the worst scores, followed by CT then CT+TH rats. The TH mean was not significantly different than Sham, primarily due to 3 female rats. **C.** In the novel object recognition test, no significant impairment was detected in injured rats compared to Sham. However, a striking difference between CT and TH females was observed, with CT-treated female rats spending significantly more time exploring the novel object than TH rats. **D.** The ability to hold a food item is impaired in all injured groups and modestly alleviated by CT and TH treatment. CT+TH treatment demonstrated significant improvement over NT. Forelimb height was measured from the floor of the apparatus then standardized for variation in animal size and position in image field by dividing this by body height. **E.** Impairment in ability to protect food item from theft, shown as the number of steals (# times food it3em stolen) was demonstrated in HIE rats and moderately improved by CT treatment. The graph in B and D, and the female graphs in A and C depict means (horizontal bars) and standard deviation (shaded bars). The graph in E and male graphs in A and C show median and interquartile ranges, as these data were not normally distributed. “f” indicates individual female rat values, “m” individual male rat values where data were pooled to satisfy the power threshold. All rats in the long-term cohort are represented in A, B D E (female rats: 6, 7, 6, 7, 7 in Sham, NT, CT, TH, and CTTH, respectively, male rats: 6, 7, 7, 6, 7). C represents all rats in the cohort except 1 CT+TH m whose video was inadvertently not recorded. NS indicates β>0.8, * represents p-values from 0.05-0.006, ** p-values = 0.005-0.0006, and *** < 0.0005.

#### NOR

The NOR test has traditionally been used to test recognition memory^60^ and has detected deficits in HIE previously ^45,61,62^. CT-treated female rats spent more time investigating the novel object than the other female groups (Fig. 4C). The measurement of memory in this NOR test truly measures remains controversial^63,64^, and here it seems more likely that visual and sensory processing were delayed due to thalamic damage in this group (Fig. 5C) rather than a gain of function in recognition memory in CT females.

**Figure 5.**
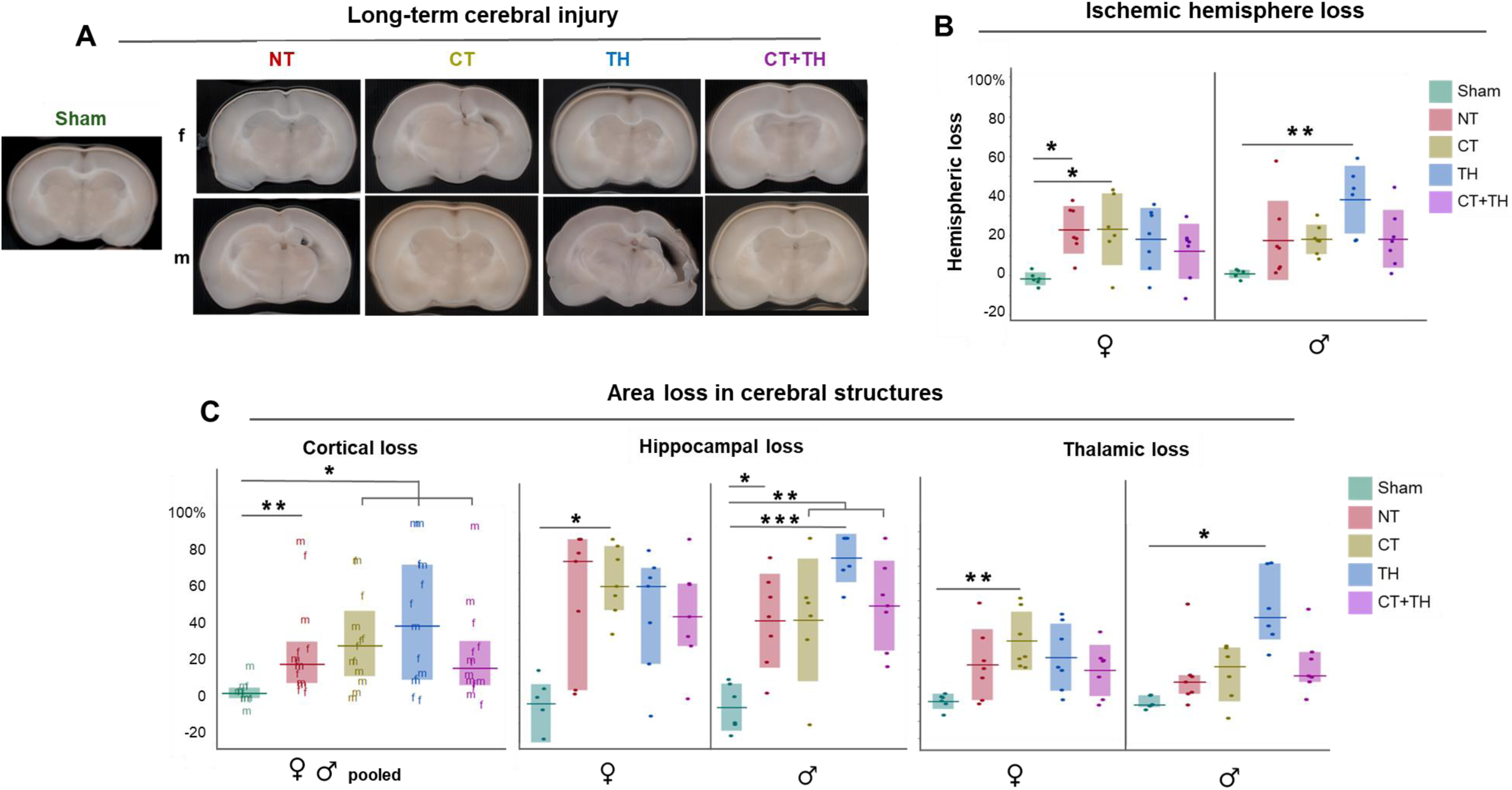
Long-term damage outcomes. **A.** Representative images of rat brains of each treatment group in the long-term harvest (66-72 dpi). Groups with HIE are delineated by sex to demonstrate the sex-specific differences. **B.** Ischemic hemisphere loss was higher in all injured rats compared to Sham, in both sexes. In females, NT and CT-treated rats showed the most tissue loss, and TH-treatment in males**. C.** Cortical, hippocampal and thalamic loss. TH-treated rats showed the most cortical loss, with TH rats demonstrating the most, followed by CT, NT, and CT+TH rats. Hippocampal loss was higher in all injured rats of both sexes compared to Sham. In females, NT rats showed the most hippocampal loss, followed by CT, TH, and CT+TH. In males TH highest, CT+TH, followed by CT and NT. Thalamic loss was higher in all injured groups relative to Sham, in both sexes. In females, CT-treated rats showed the most thalamic loss, then NT and TH, then CT+TH. In males, TH-treated rats showed the most loss, followed by CT, CT+TH, NT. Horizontal bars in the cortical, female hippocampal and male thalamic graphs represent medians, and the shaded bars interquartile range; male hippocampal and female thalamic loss graphs means and standard deviation. Scattered dots or letters represent individual rat values, with “f” representing female rat values and “m” male rat values when the sexes were pooled to meet the power threshold. *represents p-values from 0.05-0.006, ** p-values = 0.005-0.0006, and *** < 0.0005. Every rat in the long-term cohort is represented in of the measurements shown.

#### Food handling

Food handling is an activity of daily living, and the height an animal holds food items has been used previously to measure ataxia in HIE ^36,65^. Here, all injured rats held their food items lower than Sham, but CT+TH-treated held their items higher than other injured animals, and significantly more so than NT. In males, NT rats held their items the lowest, followed by TH, CT and CT+TH. Since these heights were significantly different from Sham in only NT and TH male rats (Fig. 3D), this indicates CT and CT+TH modestly improved ataxia from HIE in males more than the SOC.

#### Food protection

Protecting food is an innate behavior that draws from coordination of learning, timing, and spatial reasoning skills in rodents ^50^. It has been used to characterize brain injury, to striatum ^66^, basal ganglia, medial septum ^67^, and unilateral cortical injury ^68^. NT rats had the most food items stolen, and CT appeared to improve this ability modestly over the other treatments (Fig. 4E).

#### Long-term brain injury

Long-term brain injury was more pronounced than at short-term, as evidenced in increased tissue loss in the ischemic hemisphere (Fig. 5A) and the frequency and severity of glial scarring (Supplementary Fig. 2); most likely a reflection of chronic neuroinflammatory injury accumulated in the remainder of secondary and into tertiary phases of HIE pathology. In males, hemispheric, hippocampal and thalamic loss (Fig. 5B) were all highest with TH treatment. Taken together with the similarly poor outcomes in neuronal density and cortical and thalamic loss at 3 dpi (Fig. 2C,D), poor performance in functional testing (Fig. 4A,D), a pattern of negative effects in males by TH treatment, in our model and the conditions tested, is clear. In females, treatment with CT yielded poorer outcomes in hemispheric, hippocampal, and thalamic loss (Figs. 5B,C), and taken together with short-term and functional outcomes (Fig. 2C,D; Fig. 4A,D), CT showed the least benefit of the treatments to females.

## Discussion

In this study, we demonstrate that targeted complement modulation enhances the neuroprotective effects of TH following neonatal HI-injury. Using a clinically relevant rodent model, we show that complement-targeted therapy (CT), particularly when combined with TH (CT+TH), reduces early structural brain injury (Fig. 2), attenuates neuroinflammation (Fig. 3), and improves long-term functional and neuroanatomic outcomes (Figs 4 &5). These findings identify complement-driven immune signaling as a sustained contributor to HI-brain injury and support complement modulation as a promising adjunct to hypothermia.

A principal finding of this work is that the effects of complement activation continue beyond the acute phase of injury and contribute to secondary tissue loss. At 3 dpi, untreated animals exhibited substantial hemispheric, cortical, hippocampal, and thalamic area loss (Fig. 2), accompanied by increased apoptosis, microglial activation, astrogliosis, neuronal loss, and elevated C5aR1 expression (Fig. 3). Complement-targeted therapy, particularly when combined with hypothermia, reduced these markers of neural degeneration and inflammation, supporting a role for complement signaling in amplifying secondary injury cascades, and are consistent with prior studies implicating C5a–C5aR1 signaling in HI-injury and inflammatory neurodegeneration ^26,28,69–71^.

TH remains the SOC for neonatal HIE; however, its neuroprotective efficacy is incomplete, with nearly half of treated infants dying or surviving with adverse neurodevelopmental outcomes ^5,72,73^. Experimental and clinical evidence suggests hypothermia incompletely suppresses immune and inflammatory signaling, particularly during the subacute phase of injury ^74–76^. Our data support this concept, as hypothermia alone modestly reduced injury but did not fully attenuate inflammatory markers or prevent long-term tissue loss (Figs 2,3,5).

CT reduced inflammatory signaling even in the absence of hypothermia, and combined CT+TH treatment overall produced the most favorable outcomes across multiple measures. These findings suggest that complement signaling operates through pathways not fully addressed by temperature reduction alone and that combined targeting of metabolic and immune injury mechanisms may be necessary to achieve optimal neuroprotection.

Pronounced sex-specific differences in both injury severity and therapeutic response were observed. Males exhibited greater vulnerability to HI-insult, particularly with respect to short-term neuronal loss and structural deficits (Fig. 2). CT and hypothermia appeared to confer differential benefits by sex, with some outcomes reaching statistical significance only when sexes were analyzed separately (Figs. B, 4C, 5). In some instances, TH benefitted females but not males, with TH worsening outcomes such as hyperactivity in open field and long-term structural loss in males, when compared to untreated animals (Fig. 4A, 5). These findings align with accumulating evidence that neonatal HIE injury is sexually dimorphic, demonstrating the greater vulnerability of males to brain injury and the overall benefit of TH being possibly driven by improved outcomes in females ^36,77–79^. Failure to account for these differences may obscure therapeutic effects and limit translational success.

Functional outcomes further support the neuroprotective benefit of complement modulation, particularly in combination with hypothermia. Animals receiving CT or CT+TH demonstrated improvements in motor coordination and complex cognitive tasks such as food protection (Fig. 4) These behaviors reflect integrated sensorimotor and cognitive function and correlate with preservation of cortical, hippocampal, and thalamic structures. Prior work has shown that injury to these regions following neonatal HIE predicts long-term deficits in executive function, learning, and memory ^5,36,65,80^. Notably, functional improvements in our study were observed weeks after injury, suggesting that early modulation of complement signaling produces enduring benefits extending into later developmental stages.

Long-term neuroanatomic analysis revealed that untreated HI-injury results in progressive tissue loss extending into adolescence (Supplementary Fig. 2), consistent with ongoing tertiary injury mechanisms such as chronic inflammation, gliosis, and impaired synaptic remodeling ^81–83^. CT reduced chronic cortical, hippocampal, and thalamic atrophy (Fig. 2,5), supporting the concept that immune-mediated processes continue to influence brain remodeling long after the initial insult. These findings underscore the importance of targeting subacute and chronic inflammatory pathways in addition to acute injury mechanisms.

This study has several limitations. While the rodent model recapitulates key features of neonatal HIE, species differences in brain maturation and immune development limit direct extrapolation to human infants. However, the Vannucci model has been extensively characterized and validated over the past 40 years for neonatal HIE ^35^. Additionally, although PMX205 and C3a peptides were selected based on prior efficacy and mechanistic rationale ^31,65^, the optimal dosing, timing, and duration of complement modulation continues to be refined. Finally, while behavioral testing provides functional relevance, additional longer-term assessments into adulthood will be necessary to fully characterize cognitive and neuropsychiatric outcomes.

Despite these limitations, our findings provide evidence that complement-driven immune dysregulation contributes meaningfully to neonatal HIE injury and that targeted complement modulation enhances the neuroprotective effects of TH. By attenuating neuroinflammation, preserving neuronal integrity, and improving long-term functional outcomes, complement-targeted therapy represents a promising adjunctive strategy to current standards of care. Future studies will be required to refine therapeutic windows, define sex-specific treatment strategies, and evaluate translational potential in clinically relevant models.

## Acknowledgements

Melissa Valliere, Shelby Ridout and other members of the VHS at ODU CompMed cared for the animals. Drs. Dorela Shuboni-Mulligan and Richard Britten enriched the experiments with expert advice. Drs. Hannah Jolls, Kathryn Combo, Asna Sulaiman, Rodrigo Castro-Castenada, Amy Gaines performed end-point measurements. Mary Ann Clements embedded and sectioned the tissues.

## Data access

Data are available upon request. One author had had full access to all the data in the study and takes responsibility for its integrity and the data analysis.

## Ethics approval

This study was performed in accordance with EVMS IACUC #22-008

---

**Supplementary Figure 1.**
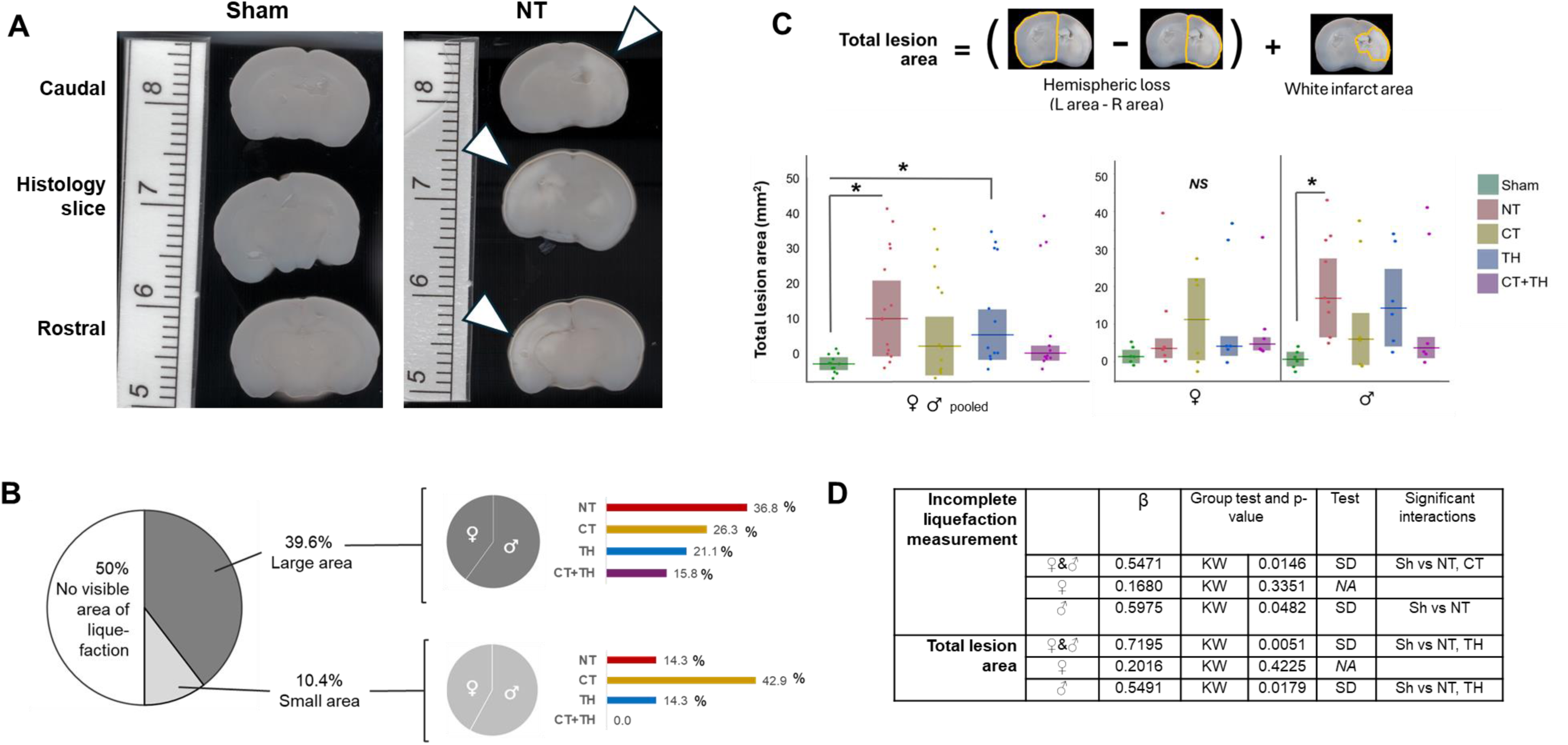
Assessments of 3 dpi brain injury including areas of incomplete liquefaction. **A.** Representative scans of uninjured (left) and injured (right) coronal brain slices. White arrows indicate the areas of incomplete liquefaction. **B**. Prevalence and severity of these areas in the cohort. 50% of injured rats showed areas of incomplete liquefaction, and of these, 39.4% showed a large area spanning the cortex, hippocampus and thalamus, that varied from completely opaque to semi-translucent, and 10.4% showed a small, less opaque area of liquefaction (approximately< 2×2 mm) confined to the upper cortex. **C.** These areas were measured in ImageJ and combined with hemispheric loss for a comprehensive measurement, but **D.** failed to reach significance. Severe and complete liquefaction was lost as dissipate into the formalin at harvest, and this tissue loss is theoretically accounted for in the hemispheric loss measurement in Fig. 2B. NS indicates group difference test did not reach significance. Every rat in the short-term cohort is represented in of the measurements shown.

**Supplementary Figure 2.**
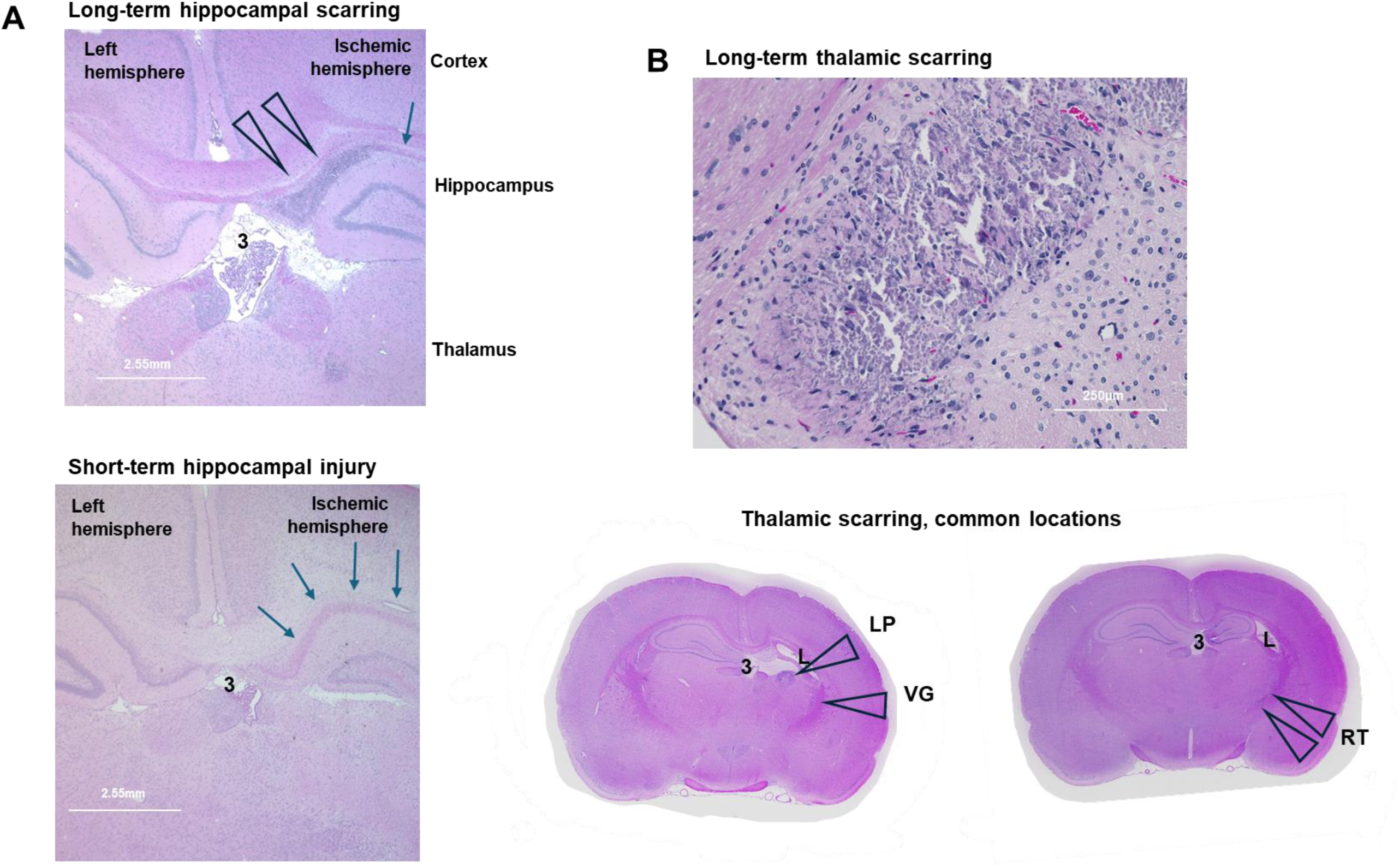
Long-term injury demonstrates hippocampal and thalamic scarring. Though rare in the short-term cohort, glial scarring was commonly detected in injured rats of the long-term cohort. **A**. Top: Representative image demonstrating the location of hippocampal scarring the long-term cohort. Bottom: analogous image of typical short-term injury, in which hippocampal scarring was rare but eosinophilic-staining neurons (a marker of neurodegeneration) were common. In both cohorts, injury to hippocampal neurons was most concentrated medially and dissipated laterally; the dentate gyrus was rarely eosinophilic but often less dense. Both images are 20x total magnification. **B.** Representative long-term thalamic scarring. Top: 200x total magnification and bottom: unmagnified but enlarged, scanned images. Thalamic scarring was present at the lateral post thalamic (LP), ventral geniculate (VG) and reticular (RT) nuclei. Throughout, hollow arrowheads highlight glial scarring, blue arrows indicate areas of eosinophilic neurons. “3” denotes the 3^rd^ ventricle, “L” the lateral ventricle. All images of H&E-stained coronal brain sections.

## References

1. Cornet MC, Kuzniewicz M, Scheffler A, et al. Perinatal Hypoxic-Ischemic Encephalopathy: Incidence Over Time Within a Modern US Birth Cohort. Pediatr Neurol. Dec 2023;149:145–150. doi:10.1016/j.pediatrneurol.2023.08.037

2. Park J, Park SH, Kim C, et al. Growth and developmental outcomes of infants with hypoxic ischemic encephalopathy. Sci Rep. Dec 28 2023;13(1):23100. doi:10.1038/s41598-023-50187-0

3. Lehtonen L, Gimeno A, Parra-Llorca A, Vento M. Early neonatal death: A challenge worldwide. Semin Fetal Neonatal Med. Jun 2017;22(3):153–160. doi:10.1016/j.siny.2017.02.006

4. Kukka AJ, Waheddoost S, Brown N, Litorp H, Wrammert J, Kc A. Incidence and outcomes of intrapartum-related neonatal encephalopathy in low-income and middle-income countries: a systematic review and meta-analysis. BMJ Glob Health. Dec 2022;7(12)doi:10.1136/bmjgh-2022-010294

5. Chakkarapani E, de Vries LS, Ferriero DM, Gunn AJ. Neonatal encephalopathy and hypoxic-ischemic encephalopathy: the state of the art. Pediatr Res. Mar 24 2025;doi:10.1038/s41390-025-03986-2

6. Mathew JL, Kaur N, Dsouza JM. Therapeutic hypothermia in neonatal hypoxic encephalopathy: A systematic review and meta-analysis. J Glob Health. 2022;12:04030. doi:10.7189/jogh.12.04030

7. Turner MJ, Dietz RM. Potential Adjuncts to Therapeutic Hypothermia to Mitigate Multiorgan Injury in Perinatal Hypoxia-Ischemia. Neoreviews. Dec 01 2023;24(12):e771–e782. doi:10.1542/neo.24-12-e771

8. Zhou KQ, Dhillon SK, Bennet L, Davidson JO, Gunn AJ. How do we reach the goal of personalized medicine for neuroprotection in neonatal hypoxic-ischemic encephalopathy? Semin Perinatol. Aug 2024;48(5):151930. doi:10.1016/j.semperi.2024.151930

9. Thayyil S, Pant S, Montaldo P, et al. Hypothermia for moderate or severe neonatal encephalopathy in low-income and middle-income countries (HELIX): a randomised controlled trial in India, Sri Lanka, and Bangladesh. Lancet Glob Health. Sep 2021;9(9):e1273–e1285. doi:10.1016/S2214-109X(21)00264-3

10. Wu YW, Comstock BA, Gonzalez FF, et al. Trial of Erythropoietin for Hypoxic-Ischemic Encephalopathy in Newborns. N Engl J Med. Jul 14 2022;387(2):148–159. doi:10.1056/NEJMoa2119660

11. Dixon BJ, Reis C, Ho WM, Tang J, Zhang JH. Neuroprotective Strategies after Neonatal Hypoxic Ischemic Encephalopathy. Int J Mol Sci. Sep 15 2015;16(9):22368–401. doi:10.3390/ijms160922368

12. Wu L, Chang E, Zhao H, Ma D. Regulated cell death in hypoxic-ischaemic encephalopathy: recent development and mechanistic overview. Cell Death Discov. Jun 11 2024;10(1):277. doi:10.1038/s41420-024-02014-2

13. Dong Q, Sun L, Peng L, et al. Expression of C5a and its receptor following spinal cord ischemia reperfusion injury in the rat. Spinal Cord. Aug 2015;53(8):581–4. doi:10.1038/sc.2015.65

14. Yuan B, Fu F, Huang S, et al. C5a/C5aR Pathway Plays a Vital Role in Brain Inflammatory Injury via Initiating Fgl-2 in Intracerebral Hemorrhage. Mol Neurobiol. 10 2017;54(8):6187–6197. doi:10.1007/s12035-016-0141-7

15. Elvington A, Atkinson C, Kulik L, et al. Pathogenic natural antibodies propagate cerebral injury following ischemic stroke in mice. J Immunol. Feb 2012;188(3):1460–8. doi:10.4049/jimmunol.1102132

16. Chan RK, Ibrahim SI, Verna N, Carroll M, Moore FD, Hechtman HB. Ischaemia-reperfusion is an event triggered by immune complexes and complement. Br J Surg. Dec 2003;90(12):1470–8. doi:10.1002/bjs.4408

17. A E, C A, H Z, et al. The alternative complement pathway propagates inflammation and injury in murine ischemic stroke. Journal of immunology (Baltimore, Md : 1950). 11/01/2012 2012;189(9)doi:10.4049/jimmunol.1201904

18. Arumugam TV, Magnus T, Woodruff TM, Proctor LM, Shiels IA, Taylor SM. Complement mediators in ischemia-reperfusion injury. Clin Chim Acta. 2006;374(1-2):33–45. doi:10.1016/j.cca.2006.06.010

19. Cowell RM, Plane JM, Silverstein FS. Complement activation contributes to hypoxic-ischemic brain injury in neonatal rats. J Neurosci. Oct 15 2003;23(28):9459–68.

20. Shah TA, Pallera HK, Kaszowski CL, Bass WT, Lattanzio FA. Therapeutic Hypothermia Inhibits the Classical Complement Pathway in a Rat Model of Neonatal Hypoxic-Ischemic Encephalopathy. Front Neurosci. 2021;15:616734. doi:10.3389/fnins.2021.616734

21. J M, F M, AL TES, VA dG, O R, AL S. Complement System in Brain Architecture and Neurodevelopmental Disorders. Frontiers in neuroscience. 02/05/2020 2020;14 doi:10.3389/fnins.2020.00023

22. Coulthard LG, Hawksworth OA, Woodruff TM. Complement: The Emerging Architect of the Developing Brain. Trends Neurosci. Jun 2018;41(6):373–384. doi:10.1016/j.tins.2018.03.009

23. Pavlovski D, Thundyil J, Monk PN, Wetsel RA, Taylor SM, Woodruff TM. Generation of complement component C5a by ischemic neurons promotes neuronal apoptosis. FASEB J. Sep 2012;26(9):3680–90. doi:10.1096/fj.11-202382

24. Farkas I, Baranyi L, Takahashi M, et al. A neuronal C5a receptor and an associated apoptotic signal transduction pathway. J Physiol. Mar 15 1998;507 ( Pt 3)(Pt 3):679–87. doi:10.1111/j.1469-7793.1998.679bs.x

25. Farkas I, Baranyi L, Liposits ZS, Yamamoto T, Okada H. Complement C5a anaphylatoxin fragment causes apoptosis in TGW neuroblastoma cells. Neuroscience. Oct 1998;86(3):903–11. doi:10.1016/s0306-4522(98)00108-0

26. Shah TA, Nejad JE, Pallera HK, et al. Therapeutic hypothermia modulates complement factor C3a and C5a levels in a rat model of hypoxic ischemic encephalopathy. Pediatr Res. Apr 2017;81(4):654–662. doi:10.1038/pr.2016.271

27. Finch AM, Wong AK, Paczkowski NJ, et al. Low-molecular-weight peptidic and cyclic antagonists of the receptor for the complement factor C5a. J Med Chem. Jun 3 1999;42(11):1965–74. doi:10.1021/jm9806594

28. Saadat A, Pallera H, Lattanzio F, et al. Structural and Functional Effects of C5aR1 Antagonism in a Rat Model of Neonatal Hypoxic-Ischemic Encephalopathy. Dev Neurosci. 2025;47(2):112–126. doi:10.1159/000539506

29. Coulthard LG, Woodruff TM. Is the complement activation product C3a a proinflammatory molecule? Re-evaluating the evidence and the myth. J Immunol. Apr 15 2015;194(8):3542–8. doi:10.4049/jimmunol.1403068

30. Moran J, Stokowska A, Walker FR, Mallard C, Hagberg H, Pekna M. Intranasal C3a treatment ameliorates cognitive impairment in a mouse model of neonatal hypoxic-ischemic brain injury. Exp Neurol. Apr 2017;290:74–84. doi:10.1016/j.expneurol.2017.01.001

31. Pekna M, Stokowska A, Pekny M. Targeting Complement C3a Receptor to Improve Outcome After Ischemic Brain Injury. Neurochem Res. Oct 2021;46(10):2626–2637. doi:10.1007/s11064-021-03419-6

32. Pozo-Rodrigálvarez A, Li Y, Stokowska A, et al. C3a Receptor Signaling Inhibits Neurodegeneration Induced by Neonatal Hypoxic-Ischemic Brain Injury. Front Immunol. 2021;12:768198. doi:10.3389/fimmu.2021.768198

33. Stokowska A, Atkins AL, Moran J, et al. Complement peptide C3a stimulates neural plasticity after experimental brain ischaemia. Brain. Feb 2017;140(2):353–369. doi:10.1093/brain/aww314

34. Ranjan AK, Gulati A. Advances in Therapies to Treat Neonatal Hypoxic-Ischemic Encephalopathy. J Clin Med. Oct 20 2023;12(20)doi:10.3390/jcm12206653

35. Vannucci SJ, Back SA. The Vannucci Model of Hypoxic-Ischemic Injury in the Neonatal Rodent: 40 years Later. Dev Neurosci. 2022;44(4-5):186–193. doi:10.1159/000523990

36. Saadat A, Blackwell A, Kaszowski C, et al. Therapeutic hypothermia demonstrates sex-dependent improvements in motor function in a rat model of neonatal hypoxic ischemic encephalopathy. Behav Brain Res. Feb 2 2023;437:114119. doi:10.1016/j.bbr.2022.114119

37. Kumar V, Lee JD, Clark RJ, Noakes PG, Taylor SM, Woodruff TM. Preclinical Pharmacokinetics of Complement C5a Receptor Antagonists PMX53 and PMX205 in Mice. ACS Omega. Feb 11 2020;5(5):2345–2354. doi:10.1021/acsomega.9b03735

38. Morán J, Stokowska A, Walker FR, Mallard C, Hagberg H, Pekna M. Intranasal C3a treatment ameliorates cognitive impairment in a mouse model of neonatal hypoxic-ischemic brain injury. Exp Neurol. 04 2017;290:74–84. doi:10.1016/j.expneurol.2017.01.001

39. Thompson SM, Berkowitz LE, Clark BJ. Behavioral and Neural Subsystems of Rodent Exploration. Learn Motiv. Feb 2018;61:3–15. doi:10.1016/j.lmot.2017.03.009

40. Eilam D, Golani I. Home base behavior of rats (Rattus norvegicus) exploring a novel environment. Behav Brain Res. Sep 1 1989;34(3):199–211. doi:10.1016/s0166-4328(89)80102-0

41. Blankenship PA, Cherep LA, Donaldson TN, et al. Otolith dysfunction alters exploratory movement in mice. Behav Brain Res. May 15 2017;325(Pt A):1–11. doi:10.1016/j.bbr.2017.02.031

42. Donaldson TN, Jennings KT, Cherep LA, et al. Antisense oligonucleotide therapy rescues disruptions in organization of exploratory movements associated with Usher syndrome type 1C in mice. Behav Brain Res. 02 15 2018;338:76–87. doi:10.1016/j.bbr.2017.10.012

43. Donaldson TN, Jennings KT, Cherep LA, et al. Progression and stop organization reveals conservation of movement organization during dark exploration across rats and mice. Behav Processes. May 2019;162:29–38. doi:10.1016/j.beproc.2019.01.003

44. Whishaw IQ, Coles BL. Varieties of paw and digit movement during spontaneous food handling in rats: postures, bimanual coordination, preferences, and the effect of forelimb cortex lesions. Behav Brain Res. May 1996;77(1-2):135–48. doi:10.1016/0166-4328(95)00209-x

45. Saadat A, Kaszowski C, Pallera H, et al. Traditional and non-traditional behavioral tests demonstrate the attenuation of cognitive deficits by therapeutic hypothermia in a rat model of neonatal hypoxic-ischemic encephalopathy. Front Behav Neurosci. 2025;19:1695435. doi:10.3389/fnbeh.2025.1695435

46. Schneider CA, Rasband WS, Eliceiri KW. NIH Image to ImageJ: 25 years of image analysis. Nat Methods. Jul 2012;9(7):671–5. doi:10.1038/nmeth.2089

47. Bogo V, Hill TA, Young RW. Comparison of accelerod and rotarod sensitivity in detecting ethanol- and acrylamide-induced performance decrement in rats: review of experimental considerations of rotating rod systems. Neurotoxicology. Dec 1981;2(4):765–87.

48. Shiotsuki H, Yoshimi K, Shimo Y, et al. A rotarod test for evaluation of motor skill learning. J Neurosci Methods. Jun 15 2010;189(2):180–5. doi:10.1016/j.jneumeth.2010.03.026

49. Sivakumaran MH, Mackenzie AK, Callan IR, Ainge JA, O’Connor AR. The Discrimination Ratio derived from Novel Object Recognition tasks as a Measure of Recognition Memory Sensitivity, not Bias. Sci Rep. Aug 1 2018;8(1):11579. doi:10.1038/s41598-018-30030-7

50. Peters MMMS. Insights from rodent food protection behaviors. Learning and Motivation. 2018;61:52–62. 10.1016/j.lmot.2017.01.004

51. Shen K, Sun J, Cao X, Zhou D, Li J. Comparison of Different Buffers for Protein Extraction from Formalin-Fixed and Paraffin-Embedded Tissue Specimens. PLoS One. 2015;10(11):e0142650. doi:10.1371/journal.pone.0142650

52. Schartz ND, Liang HY, Carvalho K, et al. C5aR1 antagonism suppresses inflammatory glial responses and alters cellular signaling in an Alzheimer’s disease mouse model. Nat Commun. Aug 15 2024;15(1):7028. doi:10.1038/s41467-024-51163-6

53. Shi Y, Jin Y, Li X, et al. C5aR1 Mediates the Progression of Inflammatory Responses in the Brain of Rats in the Early Stage after Ischemia and Reperfusion. ACS Chem Neurosci. Nov 3 2021;12(21):3994–4006. doi:10.1021/acschemneuro.1c00244

54. Van Beek J, Bernaudin M, Petit E, et al. Expression of receptors for complement anaphylatoxins C3a and C5a following permanent focal cerebral ischemia in the mouse. Exp Neurol. Jan 2000;161(1):373–82. doi:10.1006/exnr.1999.7273

55. Zhou J, Ma S, Feng D, et al. C5aR1(+) microglia exacerbate neuroinflammation and cerebral edema in acute brain injury. Neuron. Feb 4 2026;114(3):444–462 e9. doi:10.1016/j.neuron.2025.10.022

56. Alvez MB, Bergstrom S, Kenrick J, et al. A human pan-disease blood atlas of the circulating proteome. Science. Dec 18 2025;390(6779):eadx2678. doi:10.1126/science.adx2678

57. Karlsson M, Zhang C, Mear L, et al. A single-cell type transcriptomics map of human tissues. Sci Adv. Jul 2021;7(31)doi:10.1126/sciadv.abh2169

58. Woodruff TM, Crane JW, Proctor LM, et al. Therapeutic activity of C5a receptor antagonists in a rat model of neurodegeneration. FASEB J. Jul 2006;20(9):1407–17. doi:10.1096/fj.05-5814com

59. Langford-Smith A, Malinowska M, Langford-Smith KJ, et al. Hyperactive behaviour in the mouse model of mucopolysaccharidosis IIIB in the open field and home cage environments. Genes Brain Behav. Aug 2011;10(6):673–82. doi:10.1111/j.1601-183X.2011.00706.x

60. Antunes M, Biala G. The novel object recognition memory: neurobiology, test procedure, and its modifications. Cogn Process. May 2012;13(2):93–110. doi:10.1007/s10339-011-0430-z

61. Chen HR, Chen CW, Kuo YM, et al. Monocytes promote acute neuroinflammation and become pathological microglia in neonatal hypoxic-ischemic brain injury. Theranostics. 2022;12(2):512–529. doi:10.7150/thno.64033

62. Patel SD, Pierce L, Ciardiello A, et al. Therapeutic hypothermia and hypoxia-ischemia in the term-equivalent neonatal rat: characterization of a translational preclinical model. Pediatr Res. Sep 2015;78(3):264–71. doi:10.1038/pr.2015.100

63. Swiercz AP, Tsuda MC, Cameron HA. The curious interpretation of novel object recognition tests. Trends Neurosci. Apr 2025;48(4):250–256. doi:10.1016/j.tins.2025.02.003

64. Ferreira J. Rethinking the NOR Task. Lab Anim (NY). Jul 2025;54(7):176. doi:10.1038/s41684-025-01584-7

65. Saadat A, Pallera H, Lattanzio F, et al. Structural and functional effects of C5aR1 antagonism in a rat model of neonatal hypoxic-ischemic encephalopathy. Dev Neurosci. May 25 2024;doi:10.1159/000539506

66. Blankenship PA, Cheatwood JL, Wallace DG. Unilateral lesions of the dorsocentral striatum (DCS) disrupt spatial and temporal characteristics of food protection behavior. Brain Struct Funct. Aug 2017;222(6):2697–2710. doi:10.1007/s00429-017-1366-6

67. Martin MM, Winter SS, Cheatwood JL, et al. Organization of food protection behavior is differentially influenced by 192 IgG-saporin lesions of either the medial septum or the nucleus basalis magnocellularis. Brain Res. Nov 19 2008;1241:122–35. doi:10.1016/j.brainres.2008.09.018

68. Whishaw IQ. Food wrenching and dodging: use of action patterns for the analysis of sensorimotor and social behavior in the rat. J Neurosci Methods. Jun 1988;24(2):169–78. doi:10.1016/0165-0270(88)90061-1

69. Y S, Y J, X L, et al. C5aR1 Mediates the Progression of Inflammatory Responses in the Brain of Rats in the Early Stage after Ischemia and Reperfusion. ACS chemical neuroscience. 11/03/2021 2021;12(21)doi:10.1021/acschemneuro.1c00244

70. Arumugam TV, Shiels IA, Woodruff TM, Granger DN, Taylor SM. The role of the complement system in ischemia-reperfusion injury. Shock. May 2004;21(5):401–9. doi:00024382-200405000-00002 [pii]

71. K C, ND S, G B-G, et al. Modulation of C5a-C5aR1 signaling alters the dynamics of AD progression. Journal of neuroinflammation. 07/11/2022 2022;19(1)doi:10.1186/s12974-022-02539-2

72. Jacobs SE, Berg M, Hunt R, Tarnow-Mordi WO, Inder TE, Davis PG. Cooling for newborns with hypoxic ischaemic encephalopathy. Cochrane Database Syst Rev. 2013;1:Cd003311. doi:10.1002/14651858.CD003311.pub3

73. Davies A, Wassink G, Bennet L, Gunn AJ, Davidson JO. Can we further optimize therapeutic hypothermia for hypoxic-ischemic encephalopathy? Neural Regen Res. Oct 2019;14(10):1678–1683. doi:10.4103/1673-5374.257512

74. Jenkins DD, Rollins LG, Perkel JK, et al. Serum cytokines in a clinical trial of hypothermia for neonatal hypoxic-ischemic encephalopathy. Journal of Cerebral Blood Flow & Metabolism. 2012-07-18 2012;32(10):1888–1896. doi:doi:10.1038/jcbfm.2012.83

75. Zhou KQ, Bennet L, Wassink G, et al. Persistent cortical and white matter inflammation after therapeutic hypothermia for ischemia in near-term fetal sheep. J Neuroinflammation. Jun 11 2022;19(1):139. doi:10.1186/s12974-022-02499-7

76. Zhou KQ, Davidson JO. Targeting neuroinflammation after therapeutic hypothermia for perinatal hypoxic-ischemic brain injury. Neural Regen Res. Jun 2023;18(6):1261–1262. doi:10.4103/1673-5374.360174

77. Kelly LA, Branagan A, Semova G, Molloy EJ. Sex differences in neonatal brain injury and inflammation. Front Immunol. 2023;14:1243364. doi:10.3389/fimmu.2023.1243364

78. Chalak LF, Pruszynski JE, Spong CY. Sex Vulnerabilities to Hypoxia-Ischemia at Birth. JAMA Netw Open. Aug 01 2023;6(8):e2326542. doi:10.1001/jamanetworkopen.2023.26542

79. Wood TR, Gundersen JK, Falck M, et al. Variability and sex-dependence of hypothermic neuroprotection in a rat model of neonatal hypoxic-ischaemic brain injury: a single laboratory meta-analysis. Sci Rep. 07 02 2020;10(1):10833. doi:10.1038/s41598-020-67532-2

80. Glass HC, Wood TR, Comstock BA, et al. Predictors of Death or Severe Impairment in Neonates With Hypoxic-Ischemic Encephalopathy. JAMA Netw Open. Dec 02 2024;7(12):e2449188. doi:10.1001/jamanetworkopen.2024.49188

81. Levison SW, Rocha-Ferreira E, Kim BH, et al. Mechanisms of Tertiary Neurodegeneration after Neonatal Hypoxic-Ischemic Brain Damage. Pediatr Med. Aug 28 2022;5 doi:10.21037/pm-20-104

82. Fleiss B, Gressens P. Tertiary mechanisms of brain damage: a new hope for treatment of cerebral palsy? Lancet Neurol. Jun 2012;11(6):556–66. doi:10.1016/S1474-4422(12)70058-3

83. Hagberg H, Mallard C, Ferriero DM, et al. The role of inflammation in perinatal brain injury. Nat Rev Neurol. Apr 2015;11(4):192–208. doi:10.1038/nrneurol.2015.13

